# Genomic perspectives of SARS CoV-2 in liver disease patients with its clinical correlation: A single centre retrospective study

**DOI:** 10.1101/2023.02.26.530067

**Authors:** Arjun Bhugra, Reshu Agarwal, Pramod Gautam, Varun Suroliya, Ruchita Chhabra, Amit Pandey, Prince Garg, Pooja Rao, Rosmy Babu, Guresh Kumar, SM Shastry, Chhagan Bihari, Shiv Kumar Sarin, Ekta Gupta

## Abstract

**Background:** Severe Acute Respiratory Syndrome Coronavirus-2 (SARS CoV-2), is a causative agent of current global pandemic of Coronavirus disease-19 (COVID-19). Due to propagated outbreak and global vaccination drive an immense immunological selection pressure has been exerted on SARS CoV-2 leading to evolution of new variants. This study was performed to compare the mutational and clinical profile of liver disease patients infected with different variants of SARS CoV-2.

**Methodology:** This was a single-centre, retrospective, cohort study in which clinicogenomic analysis of liver disease (LD) patients infected with SARS CoV-2 was performed. Complete demographic and clinical details were retrieved from Hospital Information System (HIS). QC-threshold passed FASTA files containing sequences from COVID-19 patients (n=174) were compared with a reference genome of SARS-CoV-2 isolate named Wuhan-Hu-1 (NCBI Reference Sequence: NC_045512.2) for mutational analysis.

**Results:** Out of 232 finally analysed patients 137 (59.1%) were LD-CoV (+) and 95 (40.9%) were LD-CoV(-). LD patients with comorbidities were affected more with COVID-19 (**p=0.002**). On comparing the outcome in the terms of mortality, LD-CoV (+) had 2.29 times (**OR 2.29, CI 95%, 1.25-4.29**) higher of odds of succumbing to COVID-19 (**p=0.006**). Multivariate regression analysis revealed, abdominal distention (**p=0.05**), severe COVID-19 pneumonia (**p=0.046**) and the change in serum bilirubin levels (**p=0.005**) as well as Alkaline phosphatase (ALP) levels (**p=0.003**) to have an association with adverse outcome in LD patients with COVID-19. In Delta (22%) and Omicron (48%) groups, Spike gene harboured maximum mutations. On comparing the mutations between LD-CoV(+/D) and LD-CoV(+/O) a total of nine genes had more mutations in LD-CoV(+/O) whereas three genes had more mutations in LD-CoV(+/D).

**Conclusion:** We concluded that LD patients are more susceptible to COVID-19 as compared to a healthy adult with associated adverse clinical outcomes in terms of mortality and morbidity. Therefore this special group should be given priority while devising and introducing new vaccination and vaccination policies. The infection with different variants did not result in different outcome in our group of patients.

## 1. Introduction

Severe Acute Respiratory Syndrome Coronavirus-2 (SARS CoV-2), a causative agent of coronavirus disease 19 (COVID-19) is a novel viral pathogen which belongs to family Coronaviridae (1). COVID-19, primarily a pulmonary disease, also, affects extra-pulmonary sites with ACE-2 receptors such as liver, small intestines, heart and kidneys leading to extra-pulmonary manifestations(1,2). A few of the mechanisms which have been hypothesised for COVID-19 mediated hepatic injury are – direct hepatocytic injury impairing liver regeneration and immune responses responsible for virus clearance(3,4), increased viral replication in hepatocytes(3,5), hypoxemic vascular endothelial damage, antiviral therapy leading to drug induced liver injury (DILI) and immunoinflammatory changes(4). Involvement of liver in COVID-19 also adversely affects the clinical course and outcomes in patients with pre-existing liver disease (LD)(6–8). In general population, among patients who develop COVID-19, the prevalence of Chronic Liver Disease (CLD) patients is 2%-11%(1,9), out of which 14%-53% land up in acute hepatic dysfunction, due to severe COVID-19 pneumonia(1).

The propagated outbreak of COVID-19 and global vaccination drive have exerted an immense immunological selection pressure on SARS CoV-2. Therefore, since its first case in December 2019, five variants of concern (VOC) have evolved – Alpha (B.1.1.7), Beta (B.1.351), Gamma (P.1), Delta (B.1.617.2) and Omicron (B.1.1.529)(2,10). Within a span of two years Delta and Omicron variants have become dominant VOCs due to their increased transmissibility and decreased neutralization with vaccine or naturally acquired antibodies(2,11–14). Continues evolution of SARS CoV-2 has led to a few key mutations in Spike, ORF3a, NSP3, ORF 3 regions, associated with severe outcome in patients infected with COVID-19(15). There is very limited literature available on COVID-19 in LD patients with a detailed clinical and genomic correlation. Hence, the effect of different VOC and their key mutations on clinical course COVID-19 in LD patients is an unanswered question which needs to be addressed.

Therefore, the present study was aimed to study the clinicogenomic profile of the LD patients who were admitted with COVID-19. This was a single-centre, retrospective, cohort study in which clinicogenomic analysis of LD patients infected with SARS CoV-2 was performed at a tertiary care liver institute of North India.

## 2. Methodology

### 2.1. Study population

This was a retrospective observational study where the cases admitted with liver disease (LD) in whom COVID-19 real-time PCR testing was requested were enrolled. LD cases were defined as patients with derangement of Liver Function Tests (LFTs) for > 6 months with clinical/histological/radiological evidence of chronic changes in the liver in previous or present diagnosis. Complete demographic and clinical details were retrieved from Hospital Information System (HIS). LFTs of all the enrolled cases were recorded at two different time points: Day 0 (COVID-19 RT PCR positive) and 5^th^ day of admission/day of discharge, whichever was earlier. All LD cases were further classified into cirrhotic and non-cirrhotic.

Based on COVID-19 RT-PCR test results, all cases were divided into two groups: LD-CoV (+) and LD-CoV(-). LD-CoV (+) was further sub grouped into LD-CoV(+/D) for Delta and LD-CoV(+/O) for Omicron variant (Figure 1). COVID-19 severity was defined as per Indian Council of Medical Research Guidelines (ICMR) guidelines (https://www.icmr.gov.in/pdf/covid/techdoc/archive/COVID_Management_Algorithm_17052021.pdf).

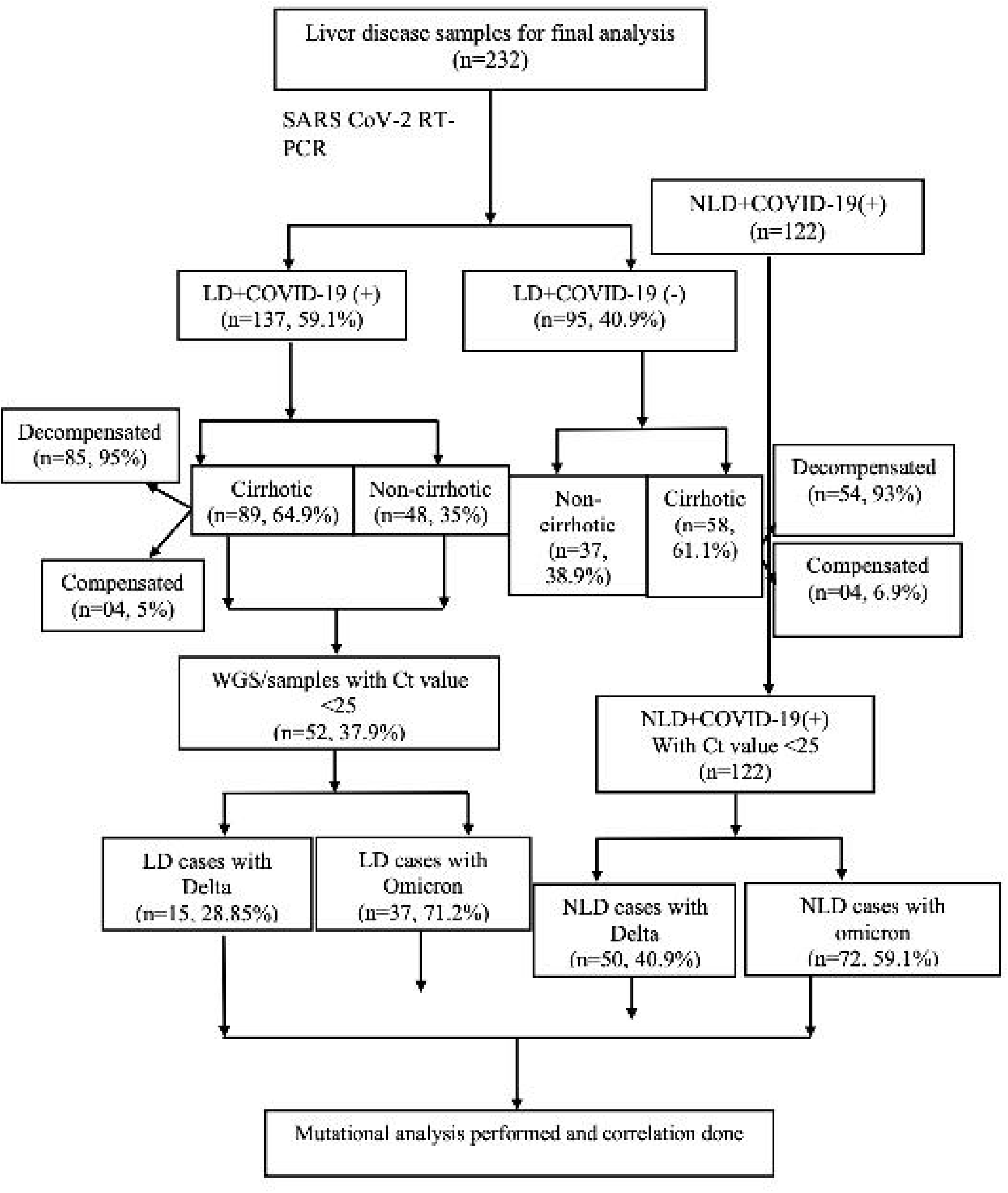

Cases with documented positive COVID-19 within 3 months from date of admission, incomplete clinical details, coinfections with HIV and other respiratory viruses and pregnant females were excluded from the analysis. For the purpose of the study admitted LD cases were enrolled corresponding to two separate COVID-19 peaks in India: 1^st^ April –30^th^ June 2021 and 1^st^ December 2021-28^th^ February 2022. The study was conducted as per the code of conduct of Declaration of Helsinki and approved by Institutional Ethics Committee (IEC) (Ethical no. IEC/2020/77/MA07).

### 2.2. SARS-CoV-2 RT-PCR test and whole genome sequencing

Total viral RNA was extracted from 300 μl of specimen (nasopharyngeal and oropharyngeal swabs in viral transport media) using Chemagic Viral DNA/RNA kit (PerkinElmer, Waltham, MA, USA) in a Chemagic 360 instrument (PerkinElmer, Waltham, MA, USA). A 10 *μ*l of the extracted viral RNA elute was further subjected to RT-PCR for the detection of SARS-CoV-2 using TRUPCR® SARS-CoV-2 realtime kit (3B Black Bio Biotech India Ltd., Bhopal, India) targeting E gene and RdRp+N gene. Sequencing of the viral isolates (Ct value <25) was done by Illumina COVIDSeq protocol on NextSeq 550 platform using ARTIC V3 Primers as per the manufacturer’s instructions. The quality check of the prepared libraries was performed using DNA high sensitivity assay kit on Bioanalyzer 2100 (Agilent Technologies, United States). The concentration of the libraries was assessed on Qubit (Thermo Fisher Scientific Inc., USA).

### 2.3. Quality control, Mapping of sequences and Lineage assignment

The raw data in the form of binary base call format (.bcl files) was generated from the NextSeq 550 instrument. These raw .bcl files were converted and de multiplexed to fastq file using bcl2fastq (Illumina, v2.20) and were aligned against the SARS-CoV-2 reference genome (NC_045512.2). The generation of a consensus genome sequence after alignment was done using the Illumina Base Space Sequence Hub. The lineage nomenclature as per the latest pango LEARN model was followed using web based or stand-alone pangolin work flows or Illumina® DRAGEN COVID Lineage App (v3.5.5) following the default parameters. In case of DRAGEN COVID Lineage tool, the minimum accepted alignment score was set to 22 and results with scores <22 were discarded. The coverage threshold and consensus sequence generation threshold were set to 20 and 90 respectively. The lineage calling target coverage which specifies the maximum number of reads with a start position overlapping any given position was set at 50. The sequencing coverage for samples was >= 30x ~ 92% and the content of non-N bases was ~ 96%.

### 2.4. Mutation profiling

For detailed mutational analysis, COVID-19 positive outpatients without any underlying liver disease during two peaks were used as control (NLD-CoV+). This group was further subdivided into NLD-CoV(+/D) for Delta and NLD-CoV(+/O) for Omicron variant (Figure 1). Finally, QC-threshold passed FASTA files containing sequences from 174 SARS-CoV-2 samples and a reference genome of SARS-CoV-2 isolate named Wuhan-Hu-1 (NCBI Reference Sequence: NC_045512.2) was used for mutational analysis.

Briefly, first multiple sequence alignment was done using MAFFT (MAFFT v7.487) with the default options. The aligned FASTA file was further refined to perform biologically relevant trimming. Finally, FastME was used for distance-based inference to create an output tree file which was used for visualization of phylogenetic tree. The annotation of nucleotide sequences was done based on NUCMER (Nucleotide Mummer) alignment tool, version 3.1 (a part of the MUMmer package). The mutations are classified according to frequency, the genomic coordinates affected and their subsequent effect on amino acid sequences. The retrieved sequences were deposited in the public repository, GISAID (https://www.gisaid.org) details of which are provided as supplementary data (Supplementary Table gisaid ids for 174 samples).

### 2.5. Statistical Analysis

Statistical Analysis was performed using SPSS software version 22. Continuous variables were summarized using mean with standard deviation or median and interquartile range (IQR) where applicable. Categorical variables were presented as percentages. To study the variance between the two groups ANOVA test was used. Univariate analysis was performed using student t-test or Wilcoxon rank sum test where applicable. Univariate and multivariate logistic regression was applied to find out the predictors in both the groups. Kaplan Meier survival analysis was also carried out. For all statistical tests, p-value less than 0.05 was considered as significant. For mutational analysis, all the statistical analysis were in R statistical environment (R version 4.1.2)

## 3. Results

Out of 267 admitted LD patients with RT-PCR requests, 232 were analysed based on inclusion criteria. Out of these 137 (59.1%) were LD-CoV (+) and remaining 95 (40.9%) were LD-CoV(-). The number of patients infected with SARS CoV-2 was slightly higher in LD-CoV (+) (59.1% vs 40.9%, p value=0.59). Both the groups were comparable as per the demographic details as mentioned in Table 1. LD patients with comorbidities especially T2DM, pre-existing lung diseases had more COVID-19 (**p=0.002**) and had higher odds of developing SARS CoV-2 infection (Table1). On comparing the outcome in the terms of mortality and mean length of stay in the hospital, LD-CoV (+) had 2.29 times (**OR 2.29, CI 95%, 1.25-4.29**) higher of odds of succumbing to COVID-19 (**p=0.006**) with a shorter mean length of stay (**p=0.002**) in the hospital.

**Table 1:** Baseline characteristics of LD-CoV (+) versus LD-CoV (-). T2DM – type 2 diabetes mellitus; HTN – hypertension, Others – Hypothyroidism, CKD, Gall stones, Renal calculus, Benign prostrate hypertrophy (BPH), Osteoporosis, hernia. Among the study population a few of the patients had both T2DM+HTN (16).

| **Demographic characteristics** | **LD-CoV (+)**<br>**(n=137,**<br>**59.05%)** | **LD-CoV (-)**<br>**(n=95,**<br>**40.94%)** | **p-value** | **OR** |
| --- | --- | --- | --- | --- |
| **Mean age, years,**<br>**(SD)** | 51.2<br>(±16.99) | 45.1<br>(±15.98) | 0.528 |  |
| **M:F** | 6.2 | 3.13 | 0.04 | 1.98<br>(1.01-3.89) |
| **Cirrhotic n(%)** | 89<br>(64.9%) | 58<br>(61%) | 0.58 | 1.18<br>(0.67-2.10) |
| Decompensated n(%) | 85 (95%) | 54 (93%) | 0.009 |  |
| Compensated n(%) | 4 (5%) | 4 (6.9%) |  |  |
| **Patients with**<br>**comorbidities n(%)** | 89<br>(64.9%) | 42<br>(44.2%) | 0.002 | 2.3<br>(1.36-3.99) |
| **T2DM n(%)** | 43<br>(48.4%) | 19<br>(45.2%) | 0.05 | 1.89<br>(1.02-3.51) |
| **HTN n(%)** | 34<br>(38.2%) | 15<br>(35.7%) | 0.06 | 1.76<br>(0.89-3.45) |
| **Pre-existing lung**<br>**diseases n(%)** | 16<br>(17.9%) | 3<br>(7.1%) | 0.043 | 3.8<br>(1.01-20.8) |
| Others | 16<br>(18%) | 14<br>(33.3%) | 0.05 | 0.44<br>(0.91-1.04) |
| Outcome |  |  |  |  |
| Mortality n (%) | 52<br>(37.9%) | 20<br>(21.1%) | 0.004 | 2.29<br>(1.25-4.29) |
| Mean length of stay (days) | 11 | 14 | 0.002 |  |
| Vaccination | n=102 | n=57 |  | 0.53<br>(0.26-1.09) |
| 2 doses | 38<br>(37.3%) | 30<br>(52.7%) | 0.067 |  |
| 1 dose or non-vaccinated | 64<br>(62.7%) | 27<br>(41.4%) |  |  |

On performing the survival analysis a Kaplan Meier graph (Figure 2) was obtained to observe the mean survival of LD patients in both the groups. Despite the higher mortality observed in LD-CoV (+) than LD-CoV (-), the mean survival in the number of days was not significantly different in two groups (**p=0.59**).

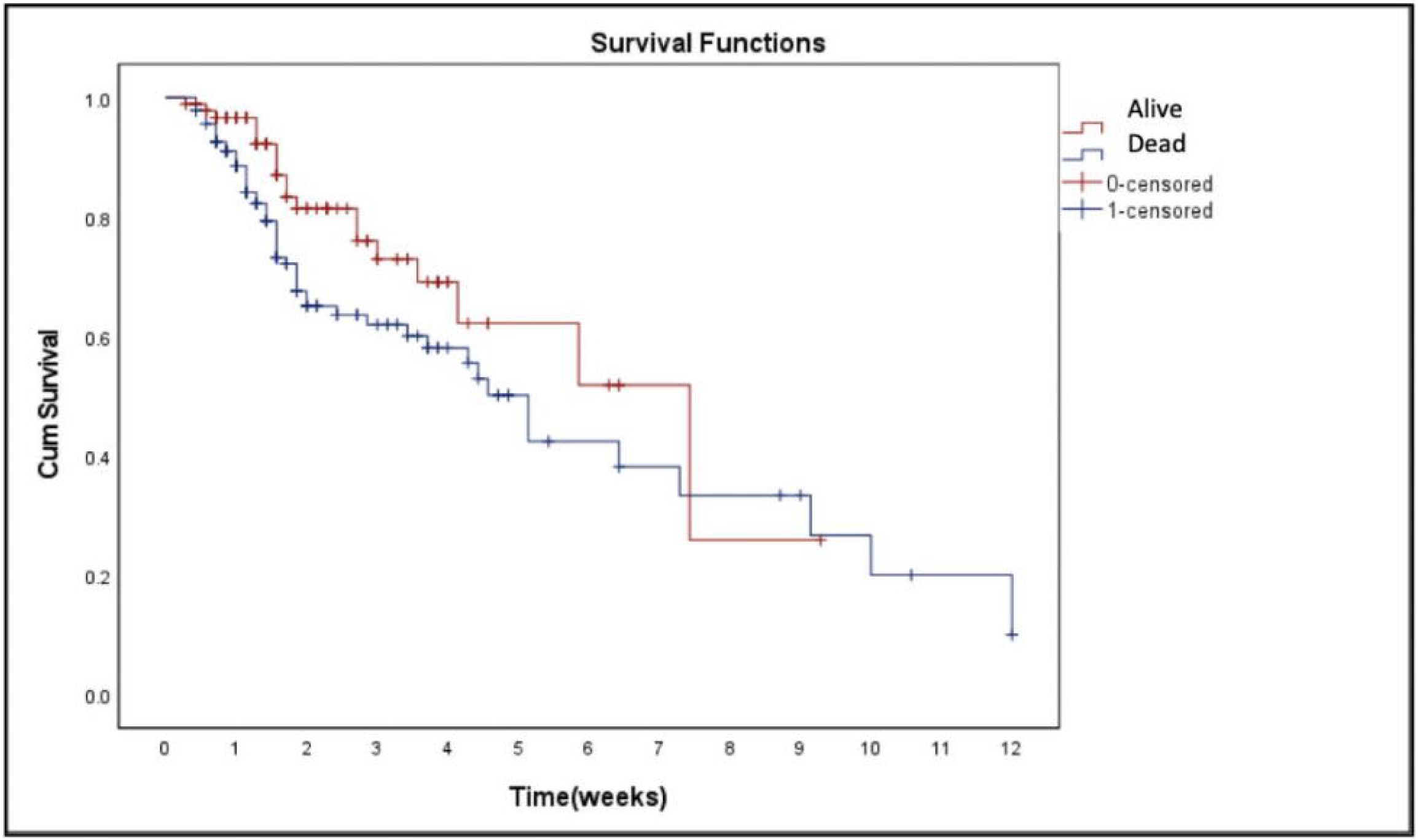

We compared the LFTs of all the enrolled cases at 2 different time points to see the effect of COVID infection on the deterioration of liver disease (Table 2). Both the groups were comparable and showed no difference in LFTs at these two time points. Though Gamma-glutamyl Transferase (GGT) levels showed a percentage increase of 13.3% in LD-CoV (+) group as compared to LD – (**p=0.07**).

**Table 2:**
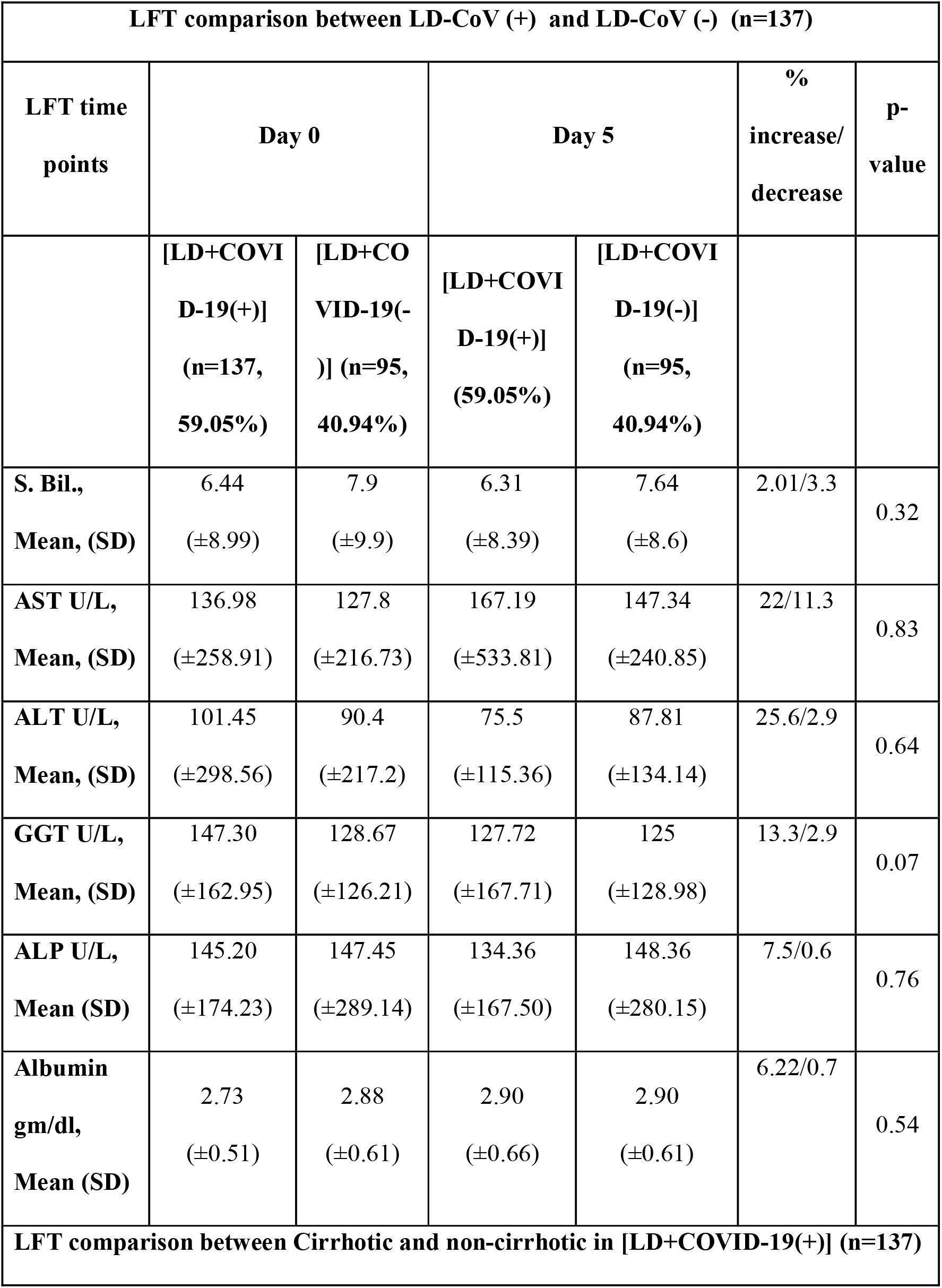
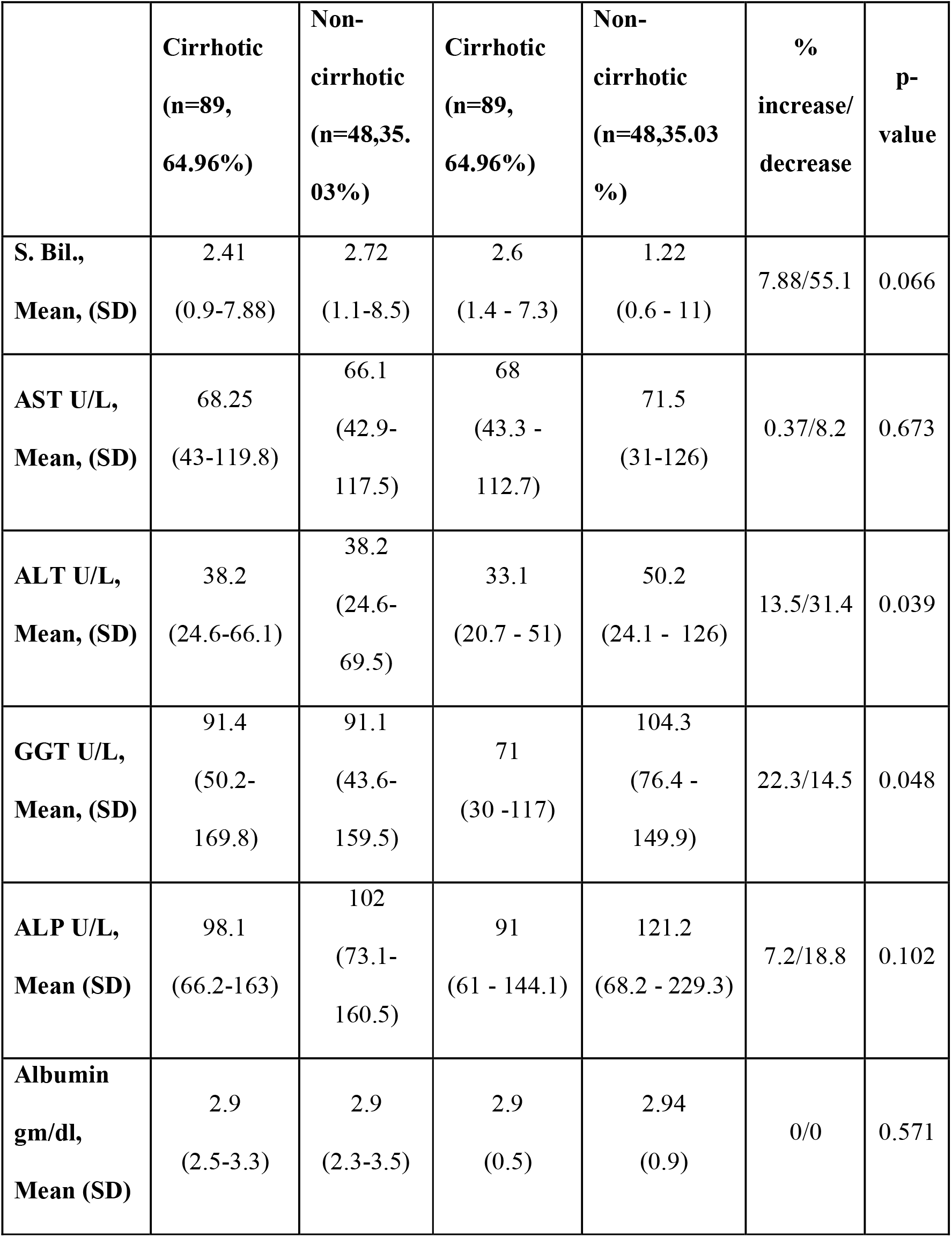
Comparison of LFT at Day 0 and Day 5 between LD patients included in the study (n=232). S. Bil. – total serum bilirubin; AST – Aspartate aminotransferase; ALT – Alanine aminotransferase; GGT – Gamma-glutamyl transferase; ALP – Alkaline phosphatase; SD – standard deviation

Multivariate regression analysis was performed taking infection with SARS CoV-2 as exposure and death as an outcome. After performing the analysis, ascites (**p=0.05**), severe COVID-19 pneumonia (**p=0.046**) and the change in serum bilirubin levels (**p=0.005**) as well as Alkaline phosphatase (ALP) levels (**p=0.003**) showed an association with adverse outcome in LD patients with COVID-19 Table S1.

### 3.1. Subgroup analysis of cirrhotic and non-cirrhotic patients in LD-CoV (+) (n=137)

Out of LD (+) group 89 were cirrhotic (64.9%) and 48 were non-cirrhotic (35.03%). Both the groups had comparable demographic characteristics as shown in Table S2. In this sub-group, non-cirrhotic LD patients (18, 37.5%) with additional comorbidity i.e. HTN showed more number of SARS CoV-2 infections than cirrhotic (18, 28.1%). Outcome in terms of mortality was slightly more in the cirrhotic group (38, 42.7%) compared to the non-cirrhotic (14, 29.2%) (p value=0.12).

On comparing the LFTs at the two different time points (Table 2), Alanine transaminase (ALT) (increase of 31.4%, **p value=0.03**) and GGT (increase of 13.5%, p value=0.048) levels showed an increase on 5^th^ day of infection in non-cirrhotic as compared to the cirrhotic group.

### 3.2. Subgroup analysis of Omicron or Delta infected patients in LD-CoV (+) (n=137)

Out of all LD-CoV (+) patients, Delta infections were recorded in 63 (45.9%) and 74 (54.01%) were infected with Omicron variant._Clinical profiles of both the groups were compared (Table S2). LD patients infected with Delta variant (25,39.7%) presented more with initial respiratory symptoms as compared to patients infected with Omicron variant (12,16.2%) (**p=0.002**). Mortality was significantly higher in Delta (38, 16.4%)than Omicron group(14, 6%) (**p<0.001**) (**OR 6.51, CI 95%, 2.83-15.23**).

### 3.3. Detection of lineages and their sub classification

Out of total 174 QC-passed and filtered sequenced samples, 65 samples were detected with Delta and 109 samples with Omicron variant. Among patients infected with Delta variant in LD-CoV (+/D) all 15 patients belonged to parent lineage i.e. B.1.617.2 and 50 patients in NLD-CoV (+/D) were infected by B.1.617.2 (47, 94%), AY. 122 (2, 4%), AY. 121 (1, 2%). Similarly among Omicron patients, 37 LD-CoV (+/O) were infected by BA.2 (32, 86.5%), BA.1.17.2 (2, 5.4%), BA.2.10 (2, 5.4%), and BA.2.37 (1, 2.7%). In NLD-CoV (+/O) group patients were infected with BA.1 (54, 75%), BA.2 (17, 24%), and BA.1.17.2 (1, 1.4%) (Figure 3).

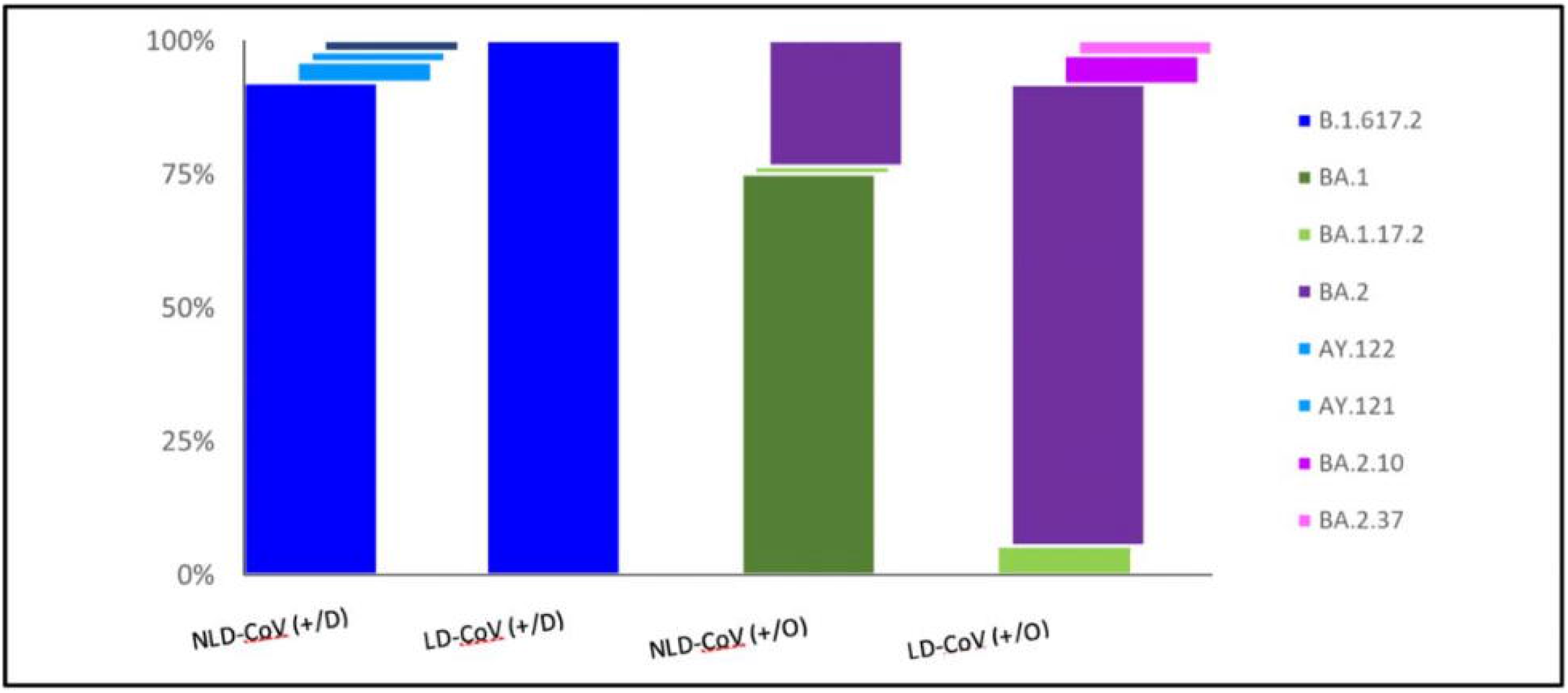

### 3.4. Mutation profiling

In Delta and Omicron groups, Spike gene harboured, ~22% and 41-48% of total mutations, respectively. In both the groups, single base substitutions were the commonest. Their numbers were – 30.2 in LD-CoV(+/D), 32.8 in NLD-CoV(+/D) and 57.1 in LD-CoV(+/O) & 49.5 NLD-CoV(+/O). Deletions were present more in Omicron (>2%) [3.9% in LD-CoV(+/O) and 5.2% in NLD-CoV(+/O)] as compared to patients in Delta (2%).

Among these 3 deletions were more prevalent in LD-CoV(+) – ORF10-CHR_END_n.29734_29759delGAGGCCACGCGGAGTACGATCGAGTG, Spike_p.L24_A27delinsS, NSP6_p.S106_F108del). Missense type of mutations were more in patients infected with Omicron (69%) than Delta (68%).

On comparing the mutations between LD-CoV(+/D) and LD-CoV(+/O) a total of nine genes (NSP4, NSP15, NS3a, NSP13, NSP5, NSP9, NSP1, NSP6, N and 3’ UTR) had more mutations in LD-CoV(+/O) whereas three genes – NSP2, NSP3 and NSP12 had more mutations in LD-CoV(+/D). Figure 4 gives an elaborate distribution of observed mutations by their type, genomic location, variation effect and putative effects.

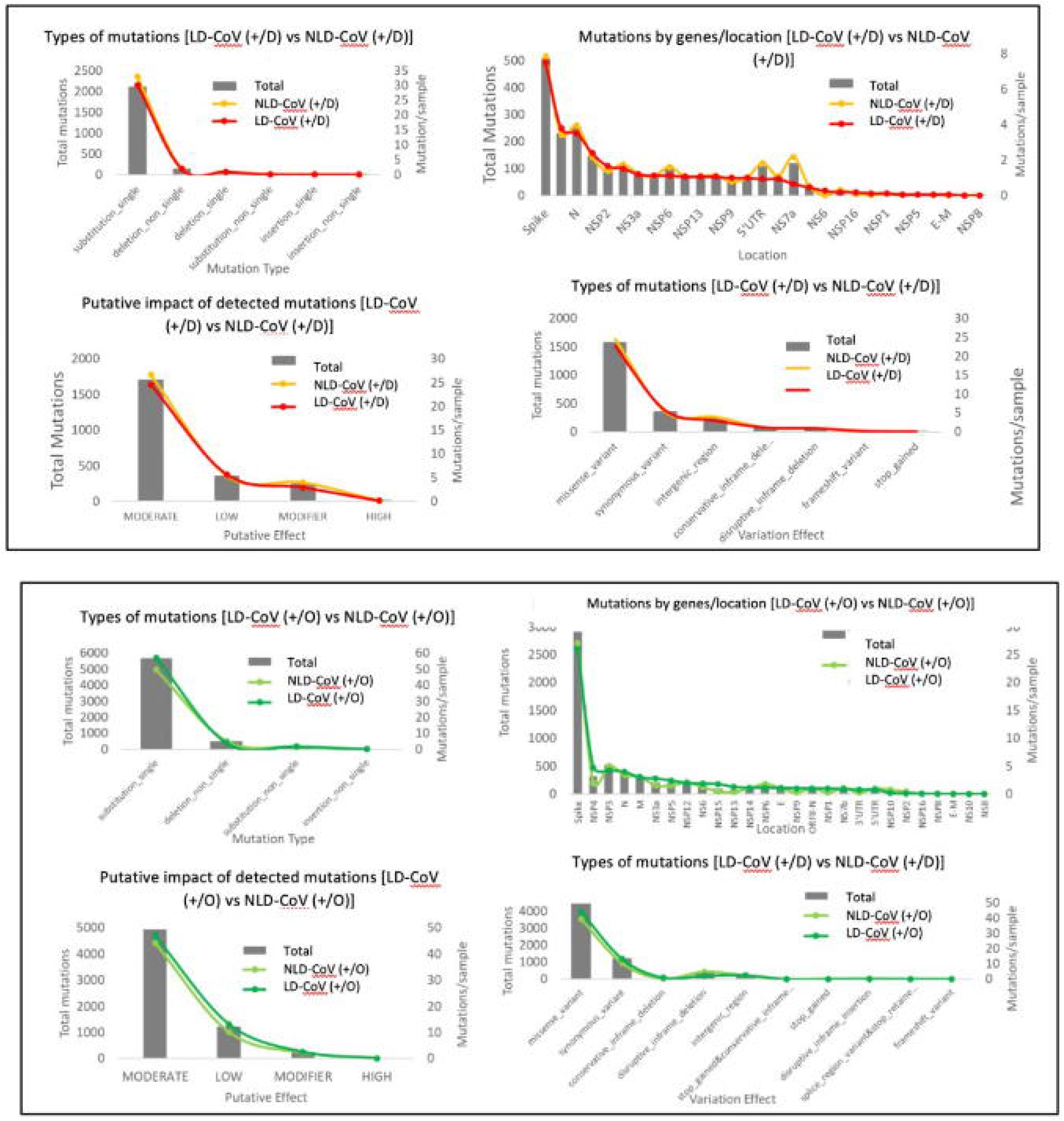

In total six genes (5’ UTR, NSP2, NSP10, NSP6, NSP3 and Spike) contained more mutations in NLD-CoV(+) compared to LD-CoV(+). Mutations between LD-CoV(+) and NLD-CoV(+) (Figure 5), were further subdivided into two categories: rare mutations (< 1% of samples) and frequent mutations (> 1% of samples). For both Delta and Omicron samples, the rare mutations were concentrated in either of the two groups but common mutations were significantly shared between the two groups (93.8% in Delta and 78.1% Omicron samples). Out of the rare mutations 60.4% to 69.3% were present in NLD-CoV(+) while 27.7% to 34.9% were present in LD-CoV(+) in Delta and Omicron data respectively.

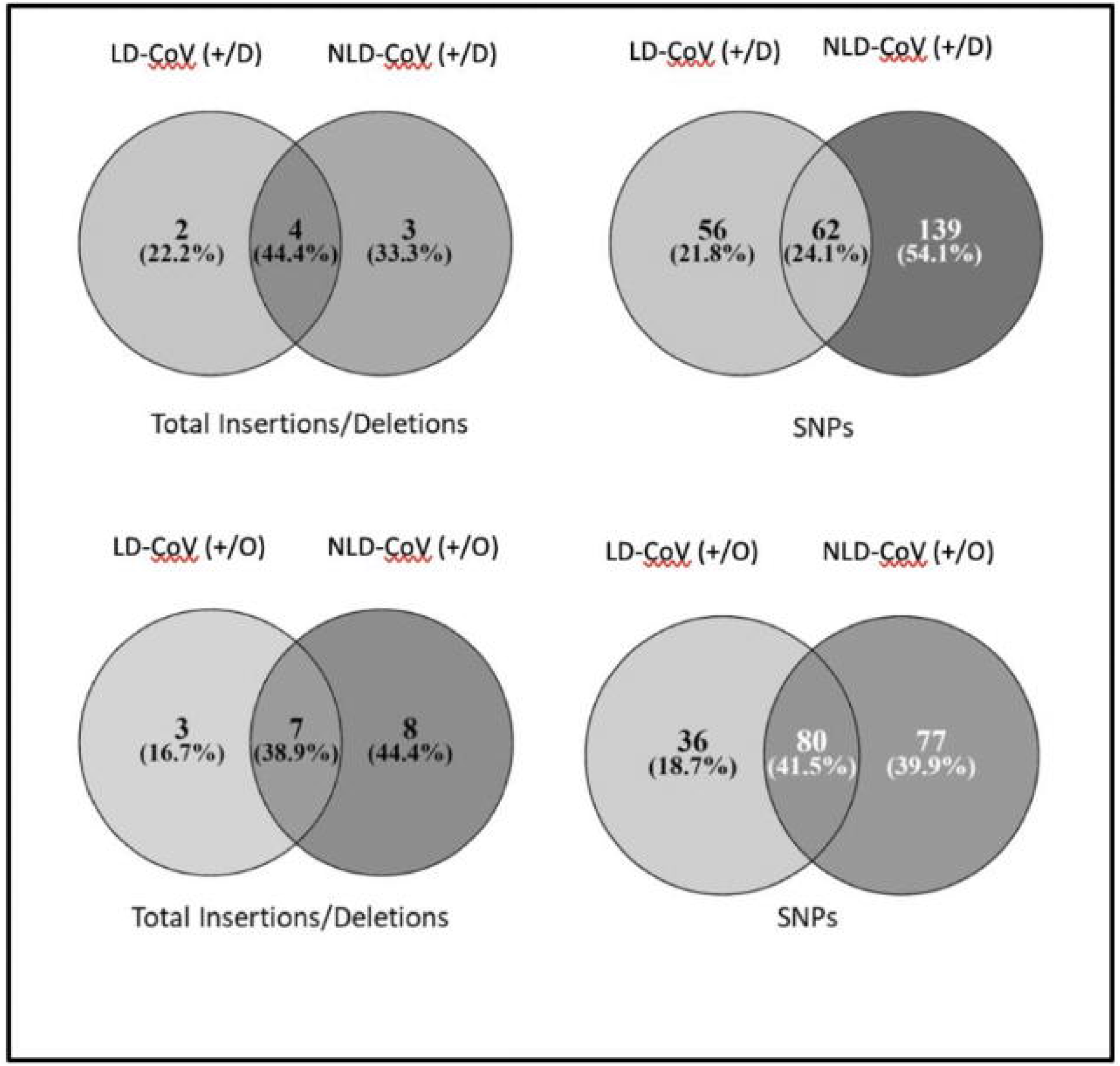

On comparing mutations between patients infected with Delta variant, LD-CoV(+/D) & NLD-CoV(+/D), 2304 total and 266 unique mutations were identified with a mean load of 33 and 36 mutations per sample respectively (**p value = 0.02**). Among these groups LD-CoV(+/D) had 13 times more mutations in NSP6 and NLD-CoV(+/D) had 3 & 2 times more mutations in NSP16 and 5’UTR region. In Omicron samples, LD-CoV(+/O) & NLD-CoV(+/O), 6419 total and 212 unique mutations were identified, with a mean load of 63 and 57 mutations per sample respectively (**p value <0.001**). Briefly, on comparing LD-CoV(+/O) & NLD-CoV(+/O), former contained more mutations in NSP4 (2.4 times), NSP13, NSP1 and 3’UTR (3.5 times). The latter contained ~5 times more mutation in NSP10 and NSP2 genes.

On analysing the putative effects of the detected mutations, ‘Moderate’ impact mutations were the most prevalent in both the groups (74-78%) with ‘Low’ impact mutation being twice more in Omicron samples (~10-13%). The percentage of shared SNPs between LD-CoV(+) and NLD-CoV(+), were 24.1% and 41.5% respectively (Figure 5). On comparing LD-CoV(+) and NLD-CoV(+), we observed that different genes showed varied frequency of mutations in Delta as well as Omicron patients in (Figure 6). On classifying shared mutation according to genomic location we observed that out of total 42 spike gene mutations in Delta samples, 9 specific to LD-CoV(+/D) and 12 shared between LD-CoV(+/D) and NLD-CoV(+/D). In Omicron group, LD-CoV(+/O) had only 1 specific mutation whereas 36 spike gene mutations were shared between LD-CoV(+/O) and NLD-CoV(+/O) (Figure 7).

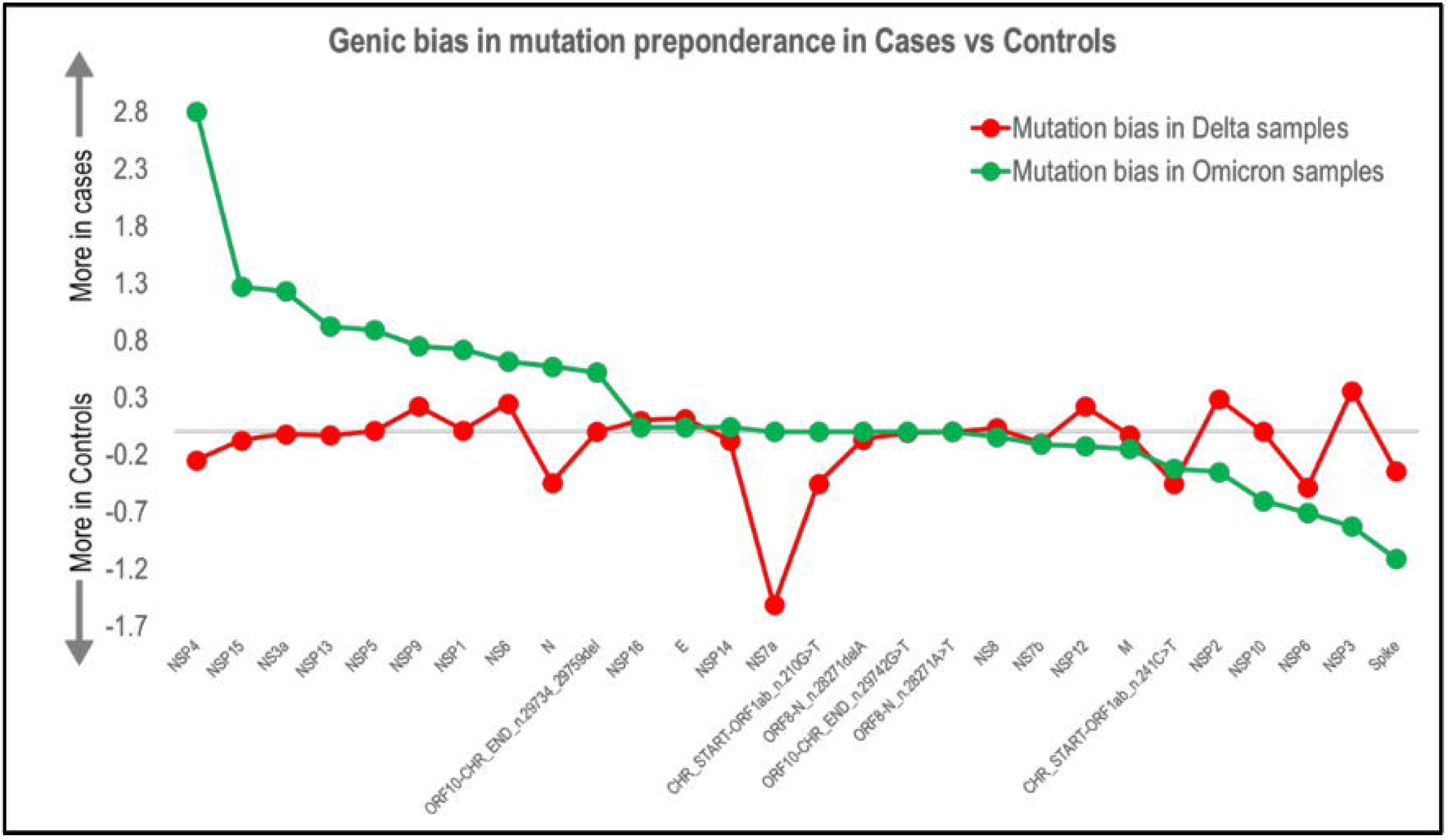

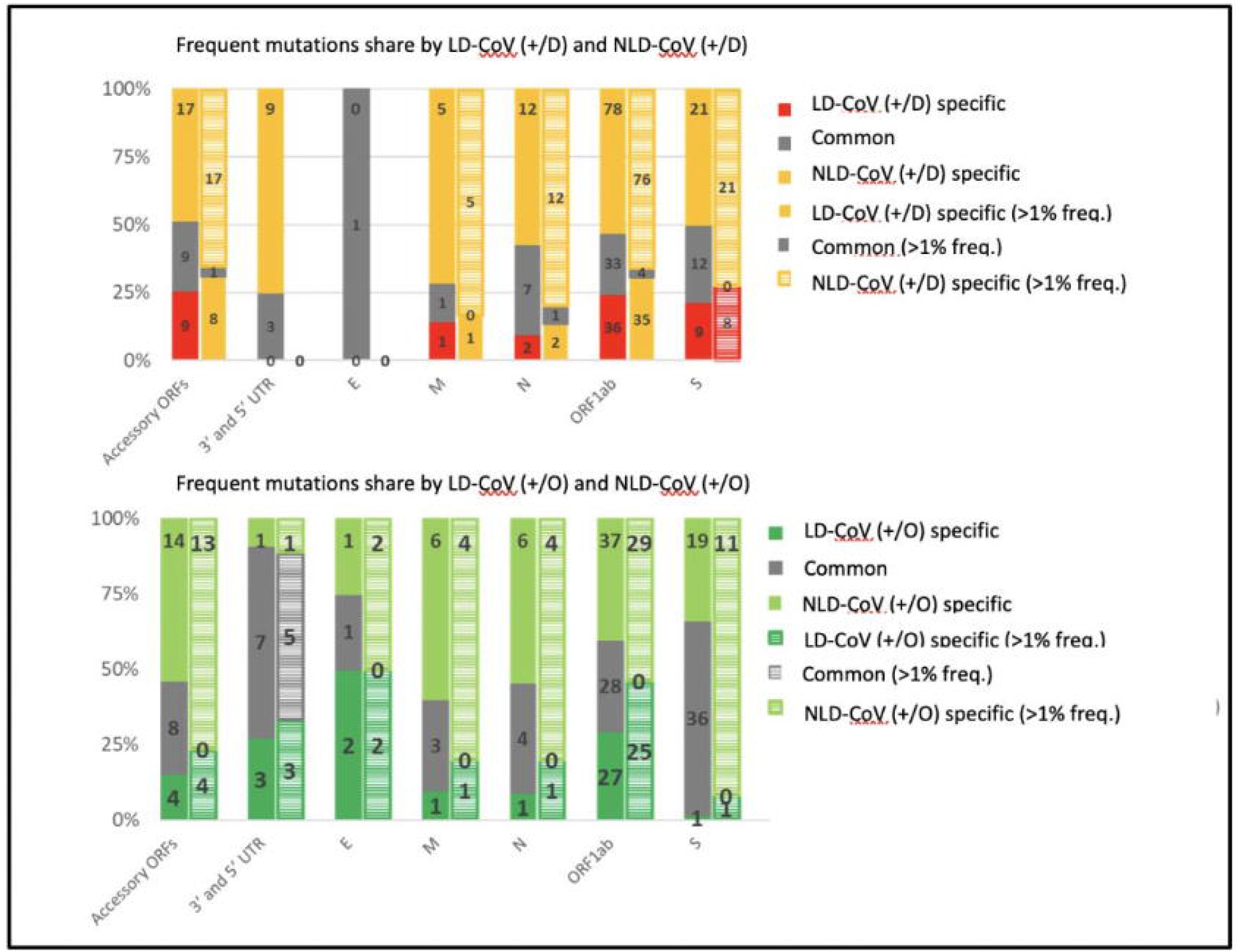

Out of 18 unique deletions, 6 were much more prevalent in NLD-CoV(+/O) (NSP6_p.L105_G107del,NSP3_p.S1265_L1266delinsI, Spike_p.G142_Y145delinsD, Spike_p.H69_V70del, Spike_p.N211_L212delinsI, Spike_p.R214_D215insEPE). Four were in Spike gene and one each in NSP3 and NSP6 gene (Figure 8).

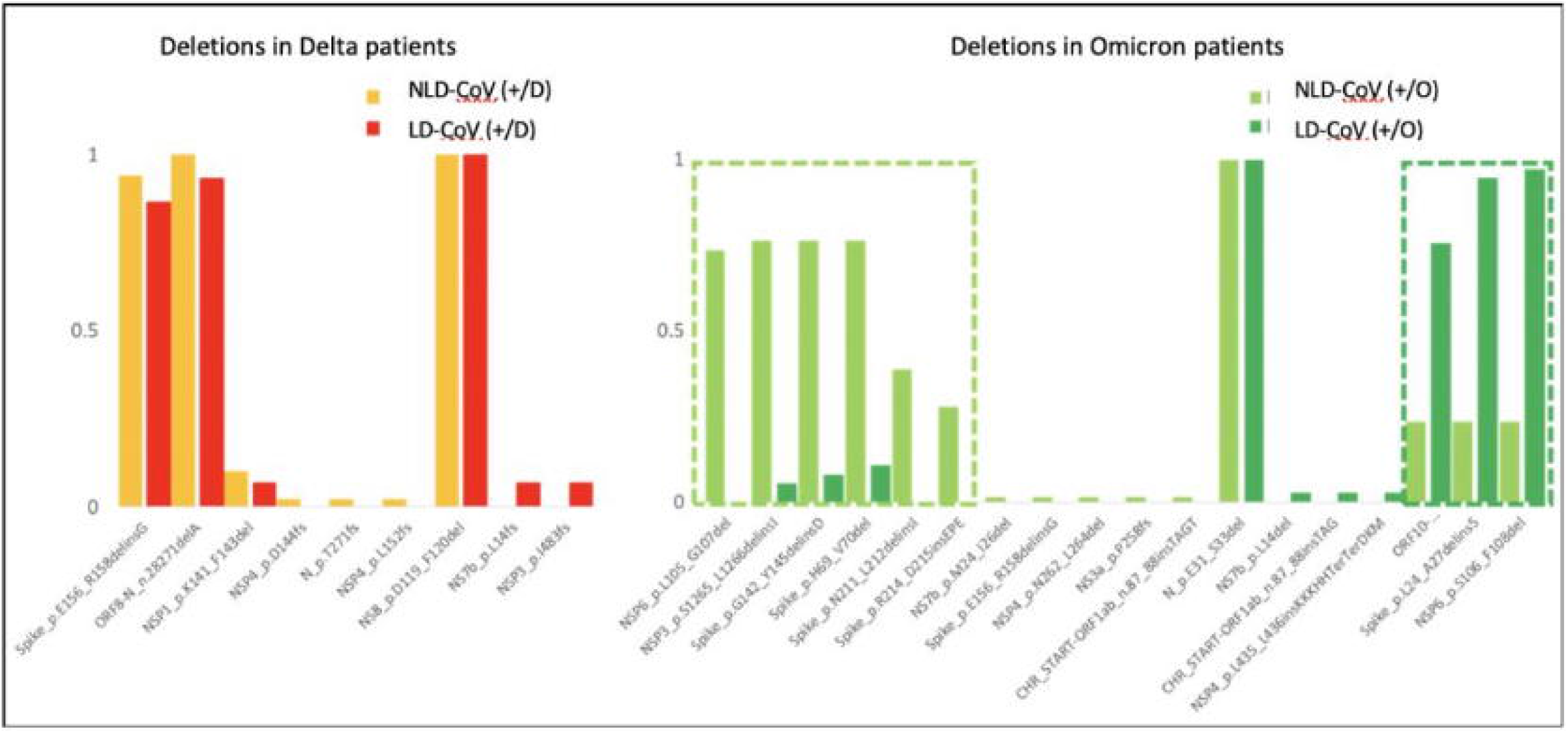

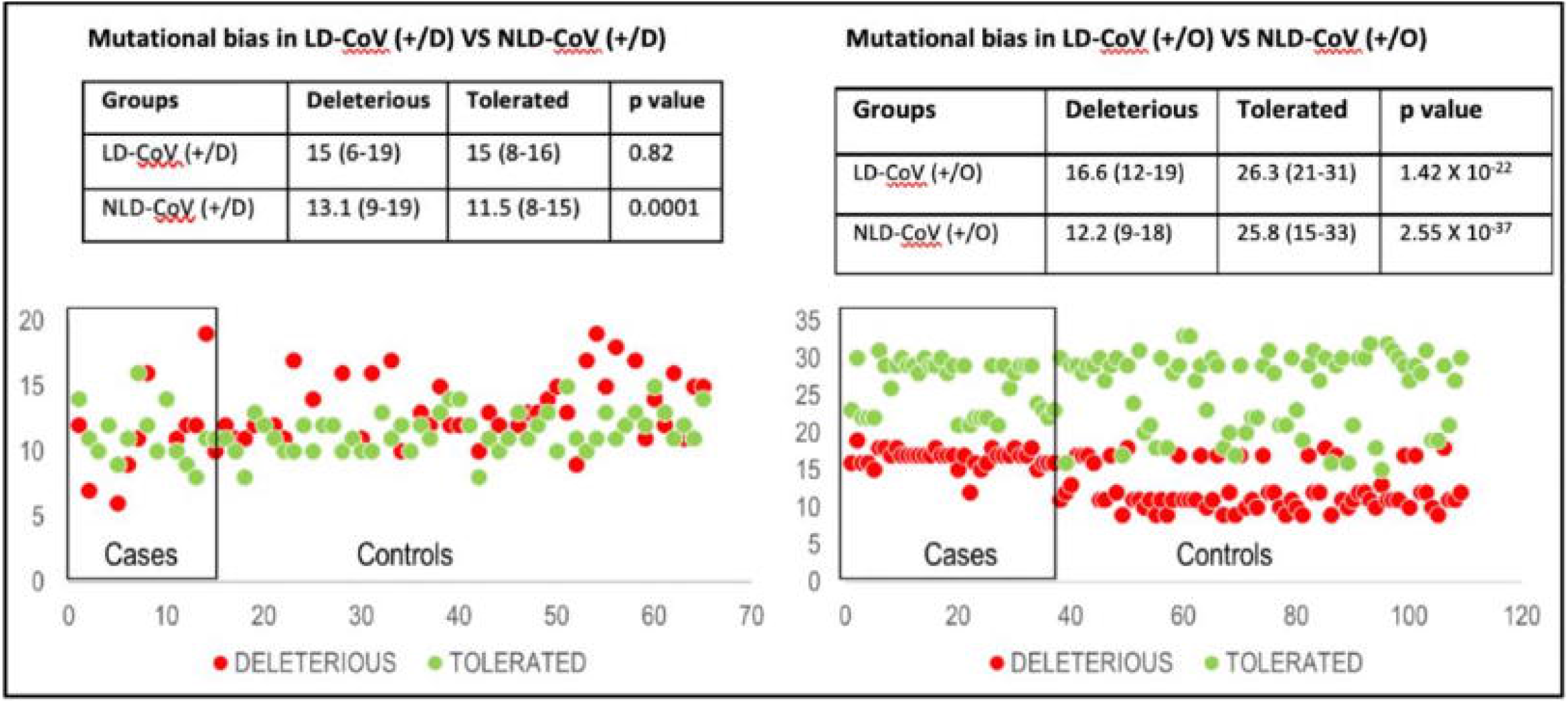

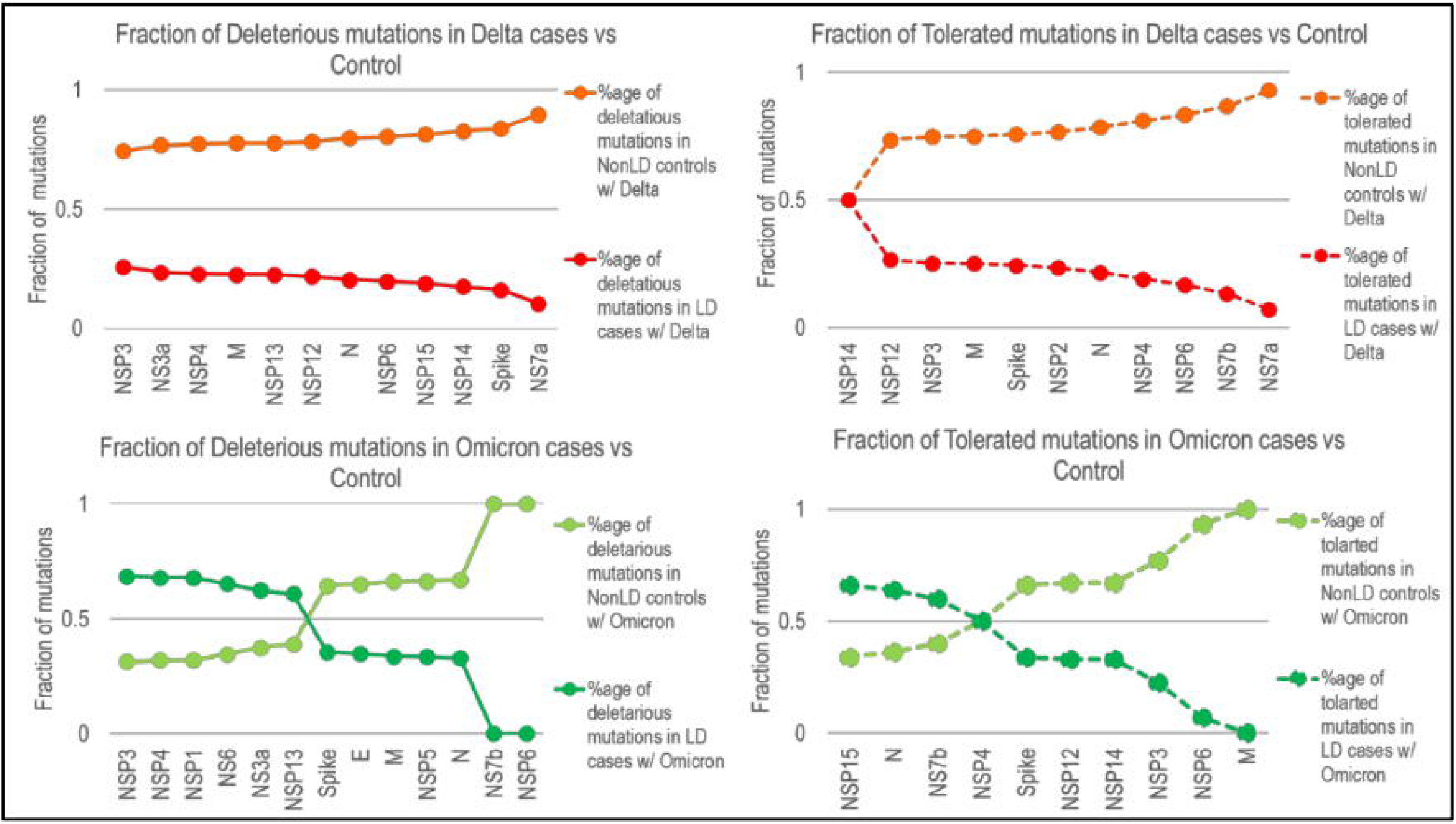

In Delta samples frequent mutations (present in >1%) were shared in miniscule fraction, while in Omicron samples only 3’ and 5’ UTR regions contained any shared mutations that were frequent too (Figure 7). Upon selecting frequent mutations it was observed that almost all of the shared mutations irrespective of their frequency (as discussed above) disappeared. N, NSP6 and NSP7a genes had more mutations in NLD-CoV(+/D) only. After analysing the deleterious and tolerated mutations, LD-CoV(+) had more deleterious mutations than NLD-CoV(+). There was significant difference between the two categories in NLD-CoV(+/D) (**p value <0.001**) with the number of deleterious mutations being significantly higher. In Omicron samples, the difference between the two categories was even more pronounced but tolerated mutations were higher in both LD-CoV(+) as well as NLD-CoV(+) (Figure 9). In Omicron samples, mutations followed two kind of distinct patterns – genes such as NSP1, NSP3, NSP4, NSP3a, NS6 and NSP13 contained higher fraction of deleterious mutations in LD-CoV(+), while genes such as E, M, N, NSP5, NSP6 and NSP7b contained higher fraction of deleterious mutations in NLD-CoV(+). Similarly, genes such as N, NSP7b and NSP15 contained higher fraction of tolerated mutations in LD-CoV(+) while genes such as S, M, NSP3, NSP6, NSP12 and NSP14 contained higher fraction of tolerated mutations in NLD-CoV(+) (Figure 10).

## 4. Discussion

To the best of our knowledge, this is one of the first study from India which has investigated and analysed correlation between clinical and genomic perspective of LD patients with COVID-19. Occurrence of COVID-19 was slightly higher in LD cases especially with other associated comorbid conditions.

In these patients, COVID-19 presents with different challenges as it is known to have a predilection and causes more severity (2,16,17). Dufour et al. have concluded that risk factors such as advancing age, decompensated cirrhosis, and presentation with gastro-intestinal (GI) symptoms have a greater risk of developing COVID-19 mediated hepatic damage(18). We had similar observations as patients with comorbidities such as T2DM, pre-existing lung diseases and HTN in LD-CoV(+) group were affected more with COVID-19. A probable reason for this can be different kinetics of SARS CoV-2 in diabetics i.e. due to elevated expression of Furin leading to increased viral hepatocyte entry, thus, helping in immune escape(19). Synergism between SARS CoV-2 and T2DM leads to upregulation of inflammatory changes and downregulation of viral interferon response leading to increased severity and mortality in patients with comorbidities(17).

On measuring the outcomes of both the groups in terms of mortality and mean length of stay in the hospital (MLS) we observed higher mortality among LD-CoV(+) as compared to LD-CoV(-). Previously, COVID-19 has been independently associated with poor clinical outcomes due to increased mortality(20). The higher risk of developing severe respiratory infection in LD patients is further complicated by secondary bacterial infections ultimately leading to hepatic complications(20,21). We did not observe any significant difference in mortality among cirrhotic and non-cirrhotic groups which was in contrast to Shalimar et al. who have observed poor outcomes in cirrhotic LD patients(22). In a different sub-group analysis which we performed, LD patients infected with Delta variant were associated with more mortality and severe disease which is in concordance with Sigal et al.(23) though the prevalence of Omicron and Delta infections in our set of patients was almost similar. We also did not observe any significant difference in different laboratory parameters which we recorded. This is in concordance with the present evidence which suggests no difference between laboratory parameters in the LD patients who suffer from Delta compared to Omicron variant, but, have increased mortality(24).

We observed that among LD-CoV(+), LFTs of non-cirrhotic patients were affected more than the cirrhotic resulting in derangement of LFTs even on the 5^th^ day of infection. Previously derangement of LFTs due to COVID-19, especially ALT and AST has been associated with higher frequency and degree of liver dysfunction in patients with or without LD(6,25). Evidence also suggests association of COVID-19 and hepatic injury resulting in persistent derangement of LFTs in patients (15%-55%) with pre-existing LD(9,26). SARS CoV-2 mediated hepatocyte damage also leads to increased severity of COVID-19 in these patients(6–8,20). In a multicentric APCOLIS study (n=228), SARS CoV-2 was shown to cause acute liver injury in non-cirrhotic patients and development of complications in approximately 50% of decompensated cirrhotic patients(6).

In this study male LD patients had higher odds of having COVID-19 as compared to females which could be owing to the differences in overall comorbidities, immune responses, or hormonal responses to COVID-19 in men and women, as described by Alwani et al(27). This is also in concordance with the previous reports on differences in clinical course of COVID-19 in both genders(16) and previous outbreaks of Severe Acute Respiratory Syndrome corona virus (SARS CoV) and Middle Eastern Respiratory Syndrome corona virus (MERS CoV)(27).

Genomic surveillance reports suggest that prolonged infections in patients with altered immune status, global mass vaccination and instillation of antiviral therapy with monoclonal antibodies puts SARS CoV-2 under constant immunological selection pressure(28–30). This has led to the development of mutations such as deletions, substitutions etc in different regions of SARS CoV-2 genome resulting in evolution of different lineages(31)

Overall observed mutations were more in samples with Omicron variant compared to Delta. We noted maximum mutations in the spike gene, Omicron (48%) more than Delta variant (22%). This was on expected lines as Omicron is one of the most divergent variant of SARS CoV-2 till date with 30 unique mutations in the spike gene region(32). (24)

Interestingly the number of unique mutations in samples with Omicron variant was lesser than Delta samples (266 as compared to 212). Genes such as NSP4, NSP15, NS3a, NSP13, NSP5, NSP9 etc. contained more mutational load in LD-CoV(+) than NLD-CoV(+). This difference in mutation numbers between LD-CoV(+/D) and NLD-CoV(+/D) was less pronounced. The shared mutations between LD-CoV(+) and NLD-CoV(+) were mostly less frequent by their occurrence which meant that the mutations specifically present either in LD-CoV(+) and NLD-CoV(+) were the frequent ones. We observed more mutation load per sample for G142D mutation in LD-CoV(+) than NLD-CoV(+). These have been previously reported from India since December 2020(33). In our analysis D614G was not associated with severe outcome as it had equal distribution of per sample load among Delta and Omicron samples. Similar kind of finding has been previously described by Long et al.(34). Substitution L452R has been linked to conferring increased infectivity in vitro, we observed to be present more in NLD-CoV(+/D) who had comparatively less mortality and severity.

Our study had certain strengths and limitations. This study is one of its kind as we compared the mutational profile of LD patients infected with Delta as well as Omicron variant. These mutations were also analysed in terms of clinical outcomes. Despite that it is a single centre study with limited number of sample size and study with much larger sample size are required to further strengthen our findings.

To conclude LD patients are more susceptible to COVID-19 as compared to a healthy adult. COVID-19 in these patients is associated with adverse clinical outcomes in terms of mortality and morbidity. Therefore this special group should be given priority while devising and introducing new vaccination and vaccination policies. Stringent droplet precautions should be followed during their ad mission especially if they have associated comorbid conditions such as T2DM, HTN and pre-existing lung diseases.

## 5. Acknowledgements

We acknowledge the Government of NCT, Delhi for facilitating the Whole Genome Sequencing Laboratory for COVID-19 at our institute. We also extend our gratitude to National Liver Disease Biobank, ILBS, New Delhi for providing Next Generation Sequencing platforms.We also thank NCDC, New Delhi and INSACOG for their unconditional support. We thank ILBS technical and administrative staff for their help and support in project implementation.

## 6. Authors Contribution

**Conceptualization:** E.G.; **Data curation:** A.B., R.C., P.G., R.A., V.S., R.B., P.G., P.R., A.P.; **Formal analysis:** P.G., G.K., A.B.; **Investigation:** V.S., P.R., P.G., R.B.; **Project administration:** S.K.S., E.G., C.B.; **Resources:** S.K.S., E.G., C.B.; **Supervision:** S.K.S., E.G., C.B.; **Visualization:** E.G., R.A., A.B.; **Writing – original draft:** A.B., R.A., P.G., V.S., G.K.; **Writing – review & editing:** S.K.S, E.G.

