## Supplemental Tables for "Genomic perspectives of SARS CoV-2 in liver disease patients with its clinical correlation: A single centre retrospective study"

| **Multivariate analysis of the study population** | | | |
| --- | --- | --- | --- |
| **Sr. No.** | **Variable** | **OR** | **p value** |
| 1 | Ascites | 3.01  (0.99-9.1) | 0.52 |
| 2 | Mild COVID-19 pneumonia | 0.96  (0.07-12.65) | 0.97 |
| 3 | Moderate COVID-19 pneumonia | 0.88  (0.4-17.88) | 0.88 |
| 4 | Severe COVID-19 pneumonia | 15.15  (1.054-217.90) | 0.04 |
| 5 | S. Bil. | 1.076  (1.023-1.13) | 0.005 |
| 6 | ALP | 0.99  (0.99-1.00) | 0.003 |

**Table S1:** Multivariate analysis of the study population taking COVID-19 as an exposure and death as an outcome. S. Bil. – Serum bilirubin, ALP – Alkaline phosphatase

| **Comparison of cirrhotic and non-cirrhotic group among LD-CoV (+) (n=137)** | | | | |
| --- | --- | --- | --- | --- |
| **Demographic characteristics** | **LD-CoV (+) Cirrhotic**  **(n=89, 64.96%)** | **LD-CoV (+)**  **Non-cirrhotic**  **(n=48,35.03%)** | **p-value** | **OR** |
| **Mean age, years,**  **Mean (SD)** | 52.61  (±16.32) | 48.85  (±18.09) | 0.22 |  |
| **Patients with comorbidities n(%)** | 57  (64%) | 32  (66.7%) | 0.75 | 0.89  (0.42-1.86) |
| T2DM n(%) | 32  (56.2%) | 11  (22.90%) | 0.09 | 1.98  (0.89-4.4) |
| HTN n(%) | 16  (28.1%) | 18  (37.5%) | 0.01 | 0.37  (0.16-0.81) |
| Pre-existing lung diseases n(%) | 14  (24.6%) | 3  (6.25%) | 0.173 | 2.8  (0.72-15.91) |
| Others | 14  (24.5%) | 2  (4.2%) | 0.03 | 4.88  (1.03-23.08) |
| **Initial respiratory symptoms at the time of presentation** | | | | |
| Fever n(%) | 28  (31.5%) | 24  (50%) | 0.03 | 0.45  (0.22-0.94) |
| Cough n(%) | 18  (20.2%) | 11  (23%) | 0.713 | 0.13  (0.36-1.99) |
| Breathing difficulty n(%) | 20  (22.5%) | 17  (35.4%) | 0.11 | 0.52  (0.24-1.14) |
| **Outcome** | | | | |
| Mortality n(%) | 38  (42.7%) | 14  (29.2%) | 0.12 | 1.81  (0.85-3.83) |
| Mean length of stay (days) | 16.7 | 16.3 | 0.54 |  |
| COVID-19 severity | | | | |
| Mild n(%) | 49  (55.1%) | 25  (52%) | 0.858 | 0.89  (0.41-1.90) |
| Moderate and severe n(%) | 40  (44.9%) | 23  (48%) |  |  |

**Table S2:** Demographics and clinical characteristics of cirrhotic and non-cirrhotic patients included in LD-CoV (+) (n=137). T2DM – type 2 diabetes mellitus; HTN – hypertension, Others – Hypothyroidism, CKD, Gall stones, Renal calculus, Benign prostrate hypertrophy (BPH), Osteoporosis, hernia. Among this group a few of the patients had both T2DM+HTN.

| **Comparison of Delta and Omicron group among LD-CoV (+) (n=137)** | | | | |
| --- | --- | --- | --- | --- |
| **Demographic characteristics** | **LD-CoV (+) Delta**  **(n=63, 45.98%)** | **LD-CoV (+) Omicron**  **(n=74, 54.01%)** | **p value** | **OR** |
| **Mean age, years (SD)** | 52.71  (±15.72) | 50.08  (±18.02) | 0.36 |  |
| **Cirrhotic n(%)** | 39  (61.9%) | 50  (67.6%) | 0.59 | 0.78  (0.36-1.67) |
| **Patients with comorbidities n(%)** | 39  (61.9%) | 50  (67.6%) | 0.48 | 0.78  (0.38-1.57) |
| T2DM n(%) | 21  (33.3%) | 23  (31.1%) | 0.77 | 1.1  (0.54-2.2) |
| HTN n(%) | 17  (27%) | 17  (23%) | 0.52 | 1.23  (0.57-2.69) |
| Pre-existing lung diseases n(%) | 8  (12.7%) | 8  (10.8%) | 0.793 | 1.2  (0.37-3.92) |
| **Initial respiratory symptoms at the time of presentation** | | | | |
| Fever n(%) | 37  (58.7%) | 15  (10.94%) | <0.001 | 5.59  (2.62-11.93) |
| Cough n(%) | 19  (13.8%) | 10  (7.2%) | 0.01 | 2.76  (1.17-6.50) |
| Breathing difficulty n(%) | 25  (39.7%) | 12  (16.2%) | 0.002 | 3.3  (1.5-7.5) |
| **Outcome** | | | | |
| Mortality n(%) | 38  (16.37%) | 14  (6.03%) | <0.001 | 6.51  (2.83-15.23) |
| Mean length of stay (days) | 11 | 10 | 0.596 |  |
| **COVID-19 severity** | | | | |
| Mild disease n(%) | 21  (33.3%) | 58  (78.4%) | <0.001 | 7.25  (3.18-16.74) |
| Moderate and severe disease (n%) | 42 (66.6%) | 16 (21.6%) |  |  |

**Table S3:** Demographics and clinical characteristics of patients infected with Delta and Omicron variant included in LD-CoV (+) (n=137). T2DM – type 2 diabetes mellitus; HTN – hypertension, Others – Hypothyroidism, CKD, Gall stones, Renal calculus, Benign prostrate hypertrophy (BPH), Osteoporosis, hernia. Among this group a few of the patients had both T2DM+HTN.
